# Transient ribosome slowdown at the decoding step triggers mRNA degradation independent of Znf598 in zebrafish

**DOI:** 10.1101/2021.05.12.443743

**Authors:** Yuichiro Mishima, Peixun Han, Seisuke Kimura, Shintaro Iwasaki

## Abstract

The control of mRNA stability plays a central role in regulating gene expression patterns. Recent studies have revealed that codon composition in the open reading frame (ORF) determines mRNA stability in multiple organisms. Based on genome-wide correlation approaches, this previously unrecognized role of the genetic code is attributable to the kinetics of the codon-decoding process by the ribosome. However, complementary experimental analysis is required to define the codon effects on mRNA stability apart from the related cotranslational mRNA decay pathways such as those triggered by aberrant ribosome stalls. In the current study, we performed a set of reporter-based analyses to define codon-mediated mRNA decay and ribosome stall-dependent mRNA decay in zebrafish embryos. Our analysis showed that the effect of codons on mRNA stability stems from the decoding process, independent of Znf598 and stall-dependent mRNA decay. We propose that codon-mediated mRNA decay is triggered by transiently slowed ribosomes engaging in a productive translation cycle in zebrafish embryos.

## Introduction

During the translation of mRNA to protein, a sequence of codons is decoded by tRNAs on the ribosome. While the translation elongation process is strictly controlled, the speed of ribosome movement passing through codons is not uniform. For example, each codon is decoded with variable efficiency due to the difference in tRNA availability (Hussmann et al., 2015; Varenne et al., 1984; Weinberg et al., 2016). Particular pairs of amino acids and codons inhibit peptide bond formation and/or decoding (Doerfel et al., 2013; Gamble et al., 2016; Schuller et al., 2017; Ude et al., 2013). The nascent polypeptide may interact with the ribosome exit tunnel and retard the ribosome (Ito and Chiba, 2013; Wilson et al., 2016). As a result, ribosomes traverse along the ORF with distinct kinetics unique to each coding sequence (Choi et al., 2018).

Nonuniform movement of the ribosome is not always harmful and is instead beneficial to the cell (Stein and Frydman, 2019). In bacteria, programmed ribosome stall by specific nascent peptides acts as a regulatory switch to control protein expression (Ito and Chiba, 2013; Wilson et al., 2016). Ribosome slowdown provides duration time for nascent chain folding and complex formation (Buhr et al., 2016; Pechmann et al., 2014; Pechmann and Frydman, 2013; Stein et al., 2019; Weinberg et al., 2016). The 5’ end of the ORF tends to contain a stretch of uncommon codons that is decoded inefficiently and optimizes the ribosome traffic right downstream of the initiation codon (Tuller et al., 2010; Weinberg et al., 2016). Hence, cells harness the nonuniform translation elongation process to control protein output.

In addition to proteome quality, ribosome movement impacts the stability of mRNAs. Transcriptome-wide analysis of mRNA half-life in budding yeast revealed a correlation between codon frequency and mRNA stability: some codons are enriched in stable mRNAs, whereas others are enriched in unstable mRNAs (Presnyak et al., 2015). Similarly, the stability of thousands of maternal mRNAs in zebrafish embryos is attributable to their codon composition (Bazzini et al., 2016; Mishima and Tomari, 2016). A correlation between codons and mRNA stability has been reported in multiple organisms based on analyses of endogenous mRNAs and ORFenome reporter mRNAs, indicating a conserved role of codons in determining mRNA stability (hereafter called codon-mediated decay) (Bazzini et al., 2016; Boël et al., 2016; Forrest et al., 2020; Harigaya and Parker, 2016; Hia et al., 2019; Narula et al., 2019; Wu et al., 2019).

As codon-mediated decay is dependent on translation, it is the ribosome that connects codons to mRNA stability (Bazzini et al., 2016; Forrest et al., 2020; Mishima and Tomari, 2016; Narula et al., 2019; Presnyak et al., 2015; Wu et al., 2019). Indeed, codon effects on mRNA stability negatively correlate with tRNA availability and ribosome density relative to A-site codons in yeast (Hanson et al., 2018). These correlations led to a model wherein codon optimality determines mRNA stability: codons that are slowly decoded due to low tRNA availability confer an mRNA-destabilizing effect, whereas codons with high tRNA availability are decoded smoothly and confer an mRNA-stabilizing effect (Presnyak et al., 2015). A subsequent study in yeast revealed that the recruitment of Not5 to the E site of the slowed ribosome and ubiquitination of ribosomal protein eS7 by Not4 are both crucial for codon-mediated decay (Buschauer et al., 2020). As Not5 and Not4 are components of the Ccr4-Not deadenylase complex, these mechanisms explain how slow ribosomes promote mRNA deadenylation and decay.

Codon-mediated decay accompanies mRNA deadenylation not only in yeast but also in zebrafish and humans (Bazzini et al., 2016; Mishima and Tomari, 2016; Presnyak et al., 2015; Webster et al., 2018; Wu et al., 2019). Furthermore, recent analyses in human cells revealed that the codon stability coefficient (CSC), ribosome density relative to the A site, and tRNA availability are correlated with each other (Forrest et al., 2020; Narula et al., 2019; Wu et al., 2019). Although tRNA availability is a complex parameter influenced by multiple factors, such as tRNA amount, tRNA aminoacylation level, and amino acid amount, these studies suggested conservation of the molecular mechanism underlying codon-mediated decay. However, genome-wide analyses and ORFeome rely on a mere correlation between mRNA stability and codon composition on a genome-wide scale and therefore lack a direct and experimental measurement of each single codon effect on mRNA stability (Bazzini et al., 2016; Forrest et al., 2020; Hanson et al., 2018; Narula et al., 2019; Presnyak et al., 2015; Wu et al., 2019).

Moreover, earlier works may include effects from related but distinct mRNA decay pathways that also act via ORFs. One such example is a translation quality control pathway that monitors the elongating ribosomes. Slowed ribosomes often induce collisions with trailing ribosomes and form specific structures called disome and trisome, a subset of which is recognized by the E3 ubiquitin ligase Hel2 in yeast. Hel2 ubiquitinates specific ribosomal proteins in the 40S subunit to initiate ribosome-associated quality control (RQC) and endonucleolytic cleavage of mRNAs via no-go decay (NGD) (Inada, 2020; Joazeiro, 2019). RQC is conserved in mammals, in which the Hel2 ortholog Znf598 acts as a collision sensor (Garzia et al., 2017; Juszkiewicz et al., 2018; Juszkiewicz and Hegde, 2017; Sundaramoorthy et al., 2017). In contrast, the occurance of NGD is controversial in vertebrates: the established RQC substrate, consecutive Lys AAA codons, does not promote apparent mRNA decay in vertebrates (Juszkiewicz and Hegde, 2017), whereas recruitment of the Znf598-4EHP-GIGYF1/2 complex on paused ribosomes promotes mRNA decay (Weber et al., 2020). Hence, it remains unclear whether the codon effects captured by genome-wide analyses in vertebrates are entirely attributable to codon-mediated decay since they may include the effect of NGD-related pathways. A simplified experimental approach that recapitulates the essence of codon-mediated decay should complement genome-wide analysis and better define the ribosome status that triggers cotranslational mRNA decay pathways.

Here, we analyze codon-mediated decay in zebrafish embryos using an artificial yet simplified reporter system. Our approach captures each single codon effect on mRNA stability in a defined context. Using this system, we evaluated the correlations between codon effects, ribosome occupancy, and the tRNA amount. We also characterized the mRNA decay pathway induced by ribosome stalls in zebrafish and analyzed its relationship to codon-mediated decay. Finally, we compared the speed of the ribosome translating mRNA-destabilizing codons to that of an aberrantly stalled ribosome. Our data provide a better definition of codon-mediated decay in zebrafish as a molecular event that occurs during slow but productive translation elongation cycles.

## Results

### Development of a codon-tag reporter system that recapitulates codon-mediated decay in zebrafish embryos

We designed an artificial codon-tag sequence to analyze the codon effects on mRNA stability in a defined sequence context (Figure 1A). This sequence contains a codon to be tested (e.g., a codon for Leu) 20 times, each separated by a codon encoding one of the 20 amino acids. The sequence was inserted at the 3’ end of the superfolder GFP (sfGFP) ORF by taking the positional effect of codon-mediated decay into account (Mishima and Tomari, 2016). We then compared the CUG and CUA Leu codon tags, whose differential effects on deadenylation were reported previously in zebrafish embryos (Mishima and Tomari, 2016). In vitro synthesized, capped and polyadenylated reporter mRNAs were injected into 1-cell stage zebrafish embryos and analyzed at two hours post fertilization (hpf) (before the maternal-to-zygotic transition [MZT]) and 6 hpf (after the MZT). Analysis of the poly(A) tail lengths by PAT assays revealed that the CUA codon-tag promoted deadenylation compared to the CUG codon-tag (Figure 1B and 1C). Consistent with the poly(A) tail status, qRT-PCR showed that the CUA codon-tag reporter was less stable than the CUG codon-tag reporter (Figure 1D). The observed difference in mRNA stability was a cotranslational effect because inhibition of translation initiation by an antisense morpholino oligonucleotide (MO) specific to the GFP ORF abolished the destabilizing effect of the CUA codon (Figure 1E). Hence, our codon-tag reporters recapitulated the translation-dependent effect of codons on mRNA deadenylation and degradation in zebrafish embryos.

**Figure 1.**
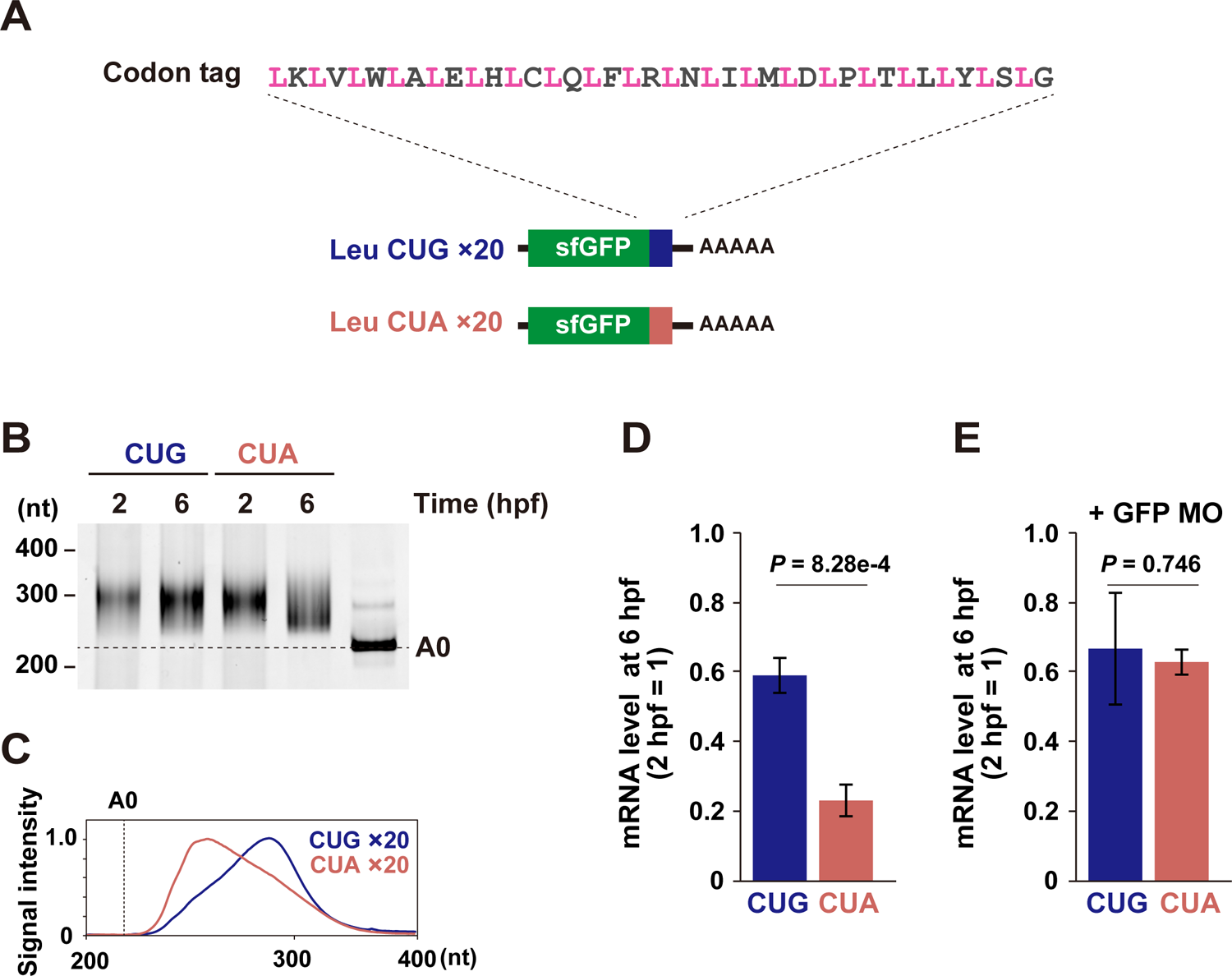
Development of a codon tag reporter system. (A) A scheme of the codon-tag reporter system. A codon to be tested (pink, in this case leucine) and one of the 20 sense codons (gray) are alternately repeated 20 times. This codon-tag sequence was inserted at the end of the sfGFP ORF. (B) Poly(A) tail analysis of codon tag reporter mRNAs at 2 and 6 hpf. The developmental stages are shown above. The lane labeled A0 shows the 3’ UTR fragment without a poly(A) tail. (C) Quantification of the PAT assay in (B). (D and E) qRT-PCR analysis of codon tag reporter mRNA levels at 6 hpf relative to 2 hpf in the absence (D) or presence (E) of translation-blocking GFP MO. Values are means ± SD of three independent experiments. P-values were calculated using two-sided Student’s t-test.

### Parallel analysis of codon effects on mRNA stability

Having validated the codon-tag reporter system with the two Leu codons, we constructed a library of codon-tag reporters to analyze multiple codon effects in parallel (hereafter called parallel analysis of codon effects; PACE). We prepared a mixture of codon-tag reporter mRNAs by plasmid cloning and in vitro transcription using this approach. We then injected a mixture of reporter mRNAs into zebrafish embryos and performed RNA sequencing (RNA-Seq) at two different time points (2 hpf and 6 hpf) to quantify the reads mapped on each tag sequence (Figure 2A). Codon-tag reads were normalized to the total reads mapped to the zebrafish genome, and the relative codon-tag amount at 6 hpf compared to 2 hpf was calculated as the effect of each codon on mRNA stability. Although four codon tags were not included due to technical limitations in library preparation and/or sequencing (UCU, UCA, GGG, and a stop codon UAG), this approach allowed us to compare the effect of 58 sense codons on mRNA stability in a defined sequence context with reasonable reproducibility (Figure 2B and S1A).

**Figure 2.**
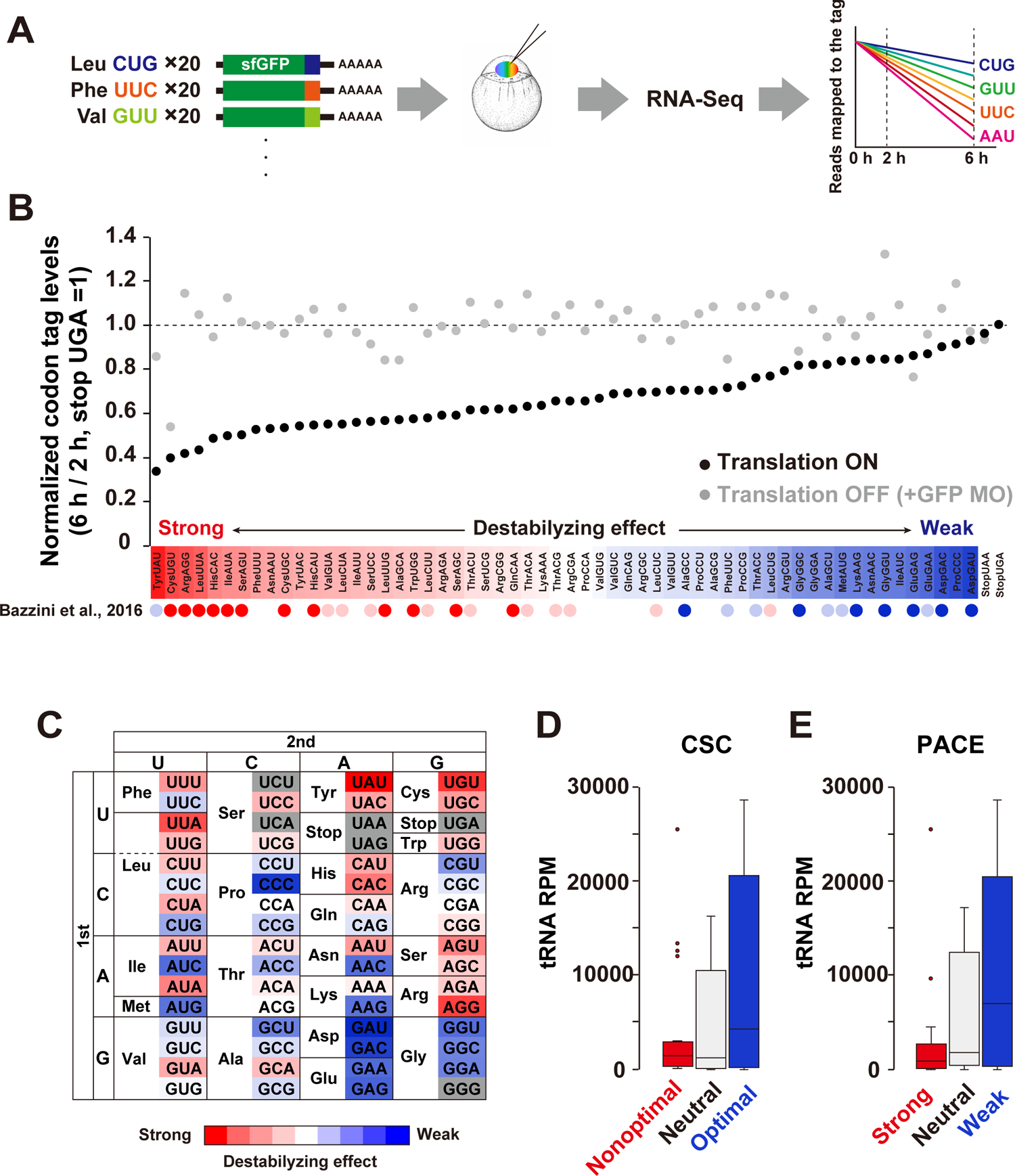
Parallel analysis of codon effects (PACE) in zebrafish embryos. (A) Scheme of parallel analysis of codon effects (PACE). Codon tag reporter mRNAs were pooled as a library and coinjected into fertilized zebrafish eggs. Embryos were collected at 2 and 6 hpf and subjected to RNA-Seq analysis. Reads mapped on each codon tag region were quantified to calculate each codon’s effect on mRNA stability. (B) Results of PACE in zebrafish embryos. Black circles show the relative stability of reporter mRNAs under normal translation. The stability of a codon-tag reporter with a UGA stop codon is set to one. Gray circles show the relative stability of reporter mRNAs under the condition in which translation initiation was inhibited by GFP MO. Averages of two biological replicates are shown. The relative effect of each codon on mRNA stability is shown as a color gradation with red (strong) to blue (weak). Codon effects estimated in Bazzini et al. 2016 are indicated below as red (nonoptimal-strong), light red (nonoptimal-weak), light blue (optimal-weak), and blue (optimal-strong) circles. (C) PACE results shown in (B) are presented as a codon table. (D) Box plot of tRNA amounts for nonoptimal (red), neutral (light gray), and optimal (blue) codons based on CSC (Bazzini et al., 2016). (E) Box plot of tRNA amounts for codons with strong (red), neutral (light gray), and weak (red) destabilizing effects based on PACE. The box represents the interquartile range (IQR), with the median indicated as a black horizontal line in the box. The whiskers represent the variation within 1.5 IQR outside the upper and lower quartiles. Outliers are shown as dots.

Several lines of evidence indicated that PACE was a valid approach for the analysis of codon-mediated decay. First, PACE detected codon-specific effects on mRNA stability. The effects were variable between synonymous codons; for example, among three synonymous codons for Ile, AUU and AUA showed a more substantial destabilizing effect than AUC (Figure 2B and 2C). A similar difference was observed in most codon boxes, although some two-codon boxes (Tyr, Cys, Asp, and Glu) and the Gly codon box showed limited variability. Second, the variable effects were dependent on translation. Inhibition of translation initiation by GFP MO stabilized almost all codon-tag reporters to a similar level, thereby reducing the difference between codons (Figure 2B and S1B). Third, PACE showed good correlations with codon frequency in the zebrafish genome (Pearson correlation coefficient *r* = 0.519; Figure S1C) and CSC (*r* = 0.689, Figure S1D), both of which were used to infer the codon effects on mRNA stability in previous genome-wide studies (Bazzini et al., 2016; Mishima and Tomari, 2016). In particular, codons classified as optimal or nonoptimal based on CSC were clearly separated in PACE (Figure 2B). We also confirmed the accuracy of PACE by qRT-PCR analysis of individual reporter mRNAs (Figure S1E and S1F).

Using CSC and tRNA sequencing, Bazzini et al. showed that nonoptimal codons tend to have fewer corresponding tRNAs than optimal codons in zebrafish (Bazzini et al., 2016) (Figure 2D). This trend was also true with the codon effects measured by PACE. When we divided codons into three groups based on the destabilizing effects in PACE (strong, neutral, and weak), we observed that codons with strong destabilizing effects tended to have fewer corresponding tRNAs (Figure 2E). Overall, we concluded that PACE successfully captured cotranslational, codon-driven effects on mRNA stability in zebrafish embryos.

### Codon effects are correlated with ribosome elongation speed and tRNA amounts

Next, we asked if the codon effects measured by PACE correlated with the ribosome elongation process. To this end, we performed ribosome footprint profiling with PACE reporters in zebrafish embryos (Han et al., 2020; Ingolia et al., 2011, 2009). We collected embryos injected with the PACE library at 4 hpf, a middle time point between 2 hpf and 6 hpf, to capture ribosome movement during the process of codon-mediated decay. Embryos were snap-frozen and treated with cycloheximide after cell lysis, which avoids aberrant effects of the compound on ribosome occupancy at codons (Hussmann et al., 2015; Weinberg et al., 2016). The obtained reads fell into 28-30 nt lengths, a hallmark of ribosome footprints (Figure S2A). Robust three-nucleotide periodicity was ensured by the information contents in the 28-30 nt fragments (Figure S2B). To analyze ribosome occupancy on codon tags, we counted the footprints with the A-site position placed on the given test codon and then normalized that by the footprints with the A site at the spacer codons (Figure 3A). This calculation allowed us to estimate the density of ribosomes waiting for decoding test codons without a complex normalization procedure. The numbers of footprints were well correlated in two replicates (Figure S2C and S2D); thus, we combined reads from two replicates to calculate the ribosome density.

**Figure 3.**
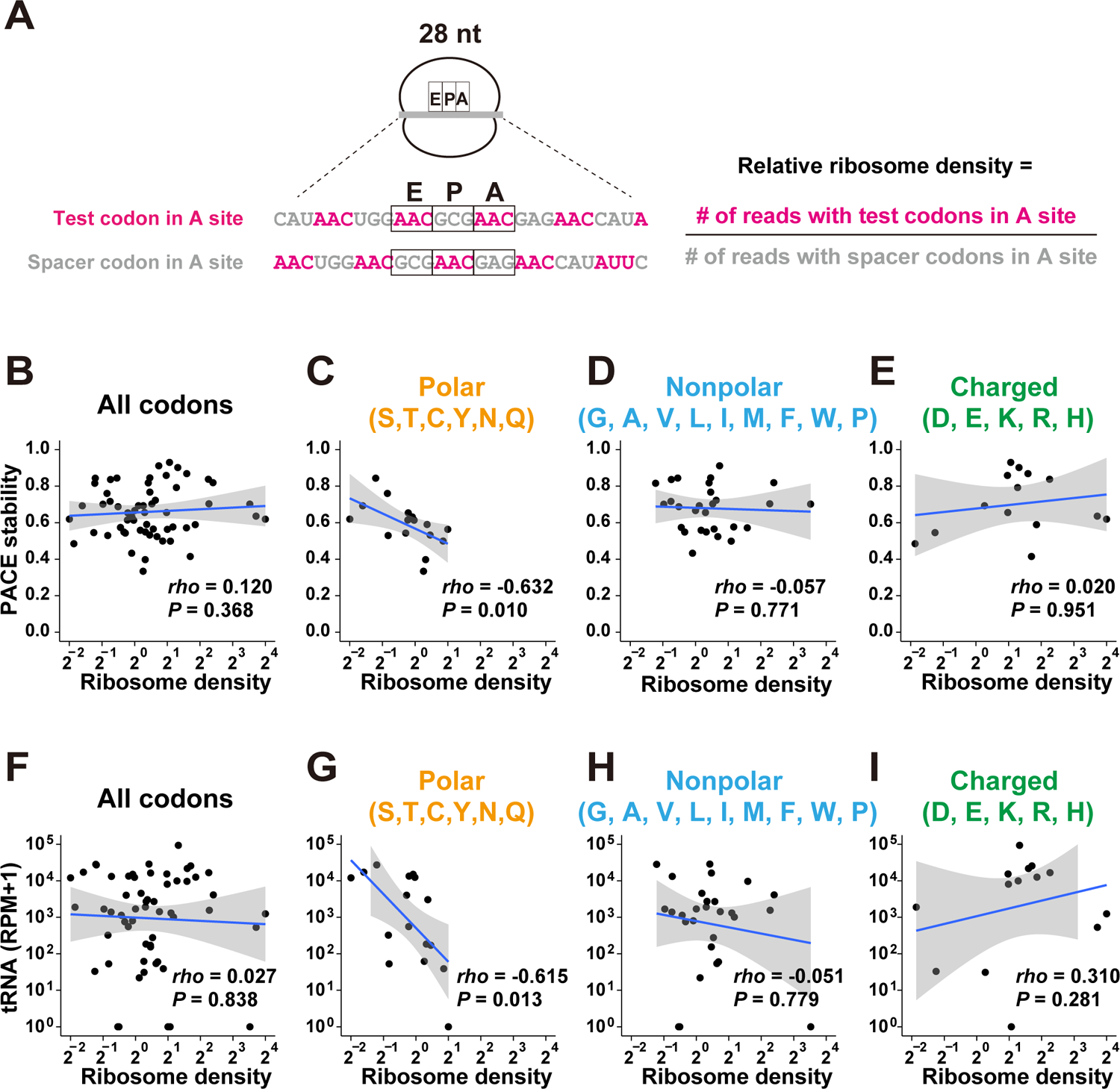
Analysis of the ribosome decoding speed at codon tag sequences. (A) Scheme of ribosome footprint analysis using PACE reporter mRNAs. An example of a 28 nt footprint is shown. The relative ribosome density of each codon at the A site was calculated by the formula presented on the right. (B-E) Scatter plots showing a correlation between relative ribosome density (28-30 nt footprints; x-axis) and the codon effects measured by PACE (y-axis). Each dot represents a single codon. Graphs for all codons (B), codons for polar amino acids (C), nonpolar amino acids (D), and charged amino acids (E) are shown. (F-I) Scatter plots showing a correlation between relative ribosome density (28-30 nt footprints; x-axis) and the tRNA amounts (y-axis, Bazzini et al., 2016). Each dot represents a single codon. Graphs for all codons (F), codons for polar amino acids (G), nonpolar amino acids (H), and charged amino acids (I) are shown. In B-I, the regression line is shown in blue and the 95% confidence interval is shown in gray. P-values were calculated using Student’s t-test.

We first analyzed a correlation between the codon effects measured by PACE and the ribosome density for all codons tested, but no clear correlation was observed (Figure 3B). Because translation of a codon tag produces a nascent peptide enriched with a particular amino acid locally, the nascent peptide within the ribosome exit tunnel may impact on the speed of ribosome traverse. Therefore, we divided the codon tags into three groups based on the characteristics of the encoded amino acid: polar, nonpolar, and charged. This classification revealed that codon tags encoding polar amino acids showed a negative correlation between the codon effects and the ribosome density: the higher the ribosome density on a given codon, the stronger the destabilization effect (Figure 3C). On the other hand, such a correlation was not observed with codons for nonpolar and charged amino acids (Figure 3D and 3E). We compared the ribosome density of tested codons to the corresponding tRNA amounts to further analyze the relationship between the ribosome density and the decoding process. Consistent with the analysis of ribosome density and the PACE results, a negative correlation between ribosome density and tRNA amounts was observed with codons for polar amino acids (Figure 3F-I). Although there was a technical limitation (see discussion), our PACE approach confirmed that codon-mediated decay is connected to the slower codon-decoding process at the A site by tRNA, at least for polar amino acid codons.

### Alteration of codon effects in response to the change in tRNA availability

Next, we modulated tRNA availability in zebrafish embryos to experimentally validate the connection between codon effects and the decoding process. The bacterial enzyme AnsB, which hydrolyzes asparagine (Asn) to aspartate (Asp), reduces aminoacylated tRNA^Asn^ and increases the duration of the ribosome on Asn codons in human cells (Loayza-Puch et al., 2016). Overexpression of AnsB in zebrafish embryos by mRNA injection caused morphological defecst at the shield stage (Figure 4A). Analysis of charged tRNA levels in these embryos by OXOPAP (Gaston et al., 2008) followed by qRT-PCR revealed that aminoacylated tRNA^Asn^_GUU_ was specifically reduced in the presence of AnsB (Figure 4B). We then examined whether the stable Asn AAC codon tag reporter mRNA was subject to codon-mediated decay in the presence of AnsB. We found that the AAC codon tag reporter mRNA was less stable in AnsB-expressing embryos than in control embryos (Figure 4C). In contrast, the stability of the unrelated codon tag reporter mRNA for Leu CUG was not affected by AnsB. The PAT assay revealed that deadenylation of the AAC codon tag reporter mRNA was promoted in AnsB-expressing embryos (Figure 4D and 4E), whereas the poly(A) tail status of the CUG codon tag reporter mRNA was not affected (Figure 4F and 4G). Hence, experimental modulation of a particular tRNA availability (tRNA^Asn^_GUU_) altered the effect of the corresponding codon (AAC, which is decoded by tRNA^Asn^_GUU_). These results support the idea that tRNA availability is a major determinant of codon-mediated deadenylation and mRNA degradation in zebrafish embryos.

**Figure 4.**
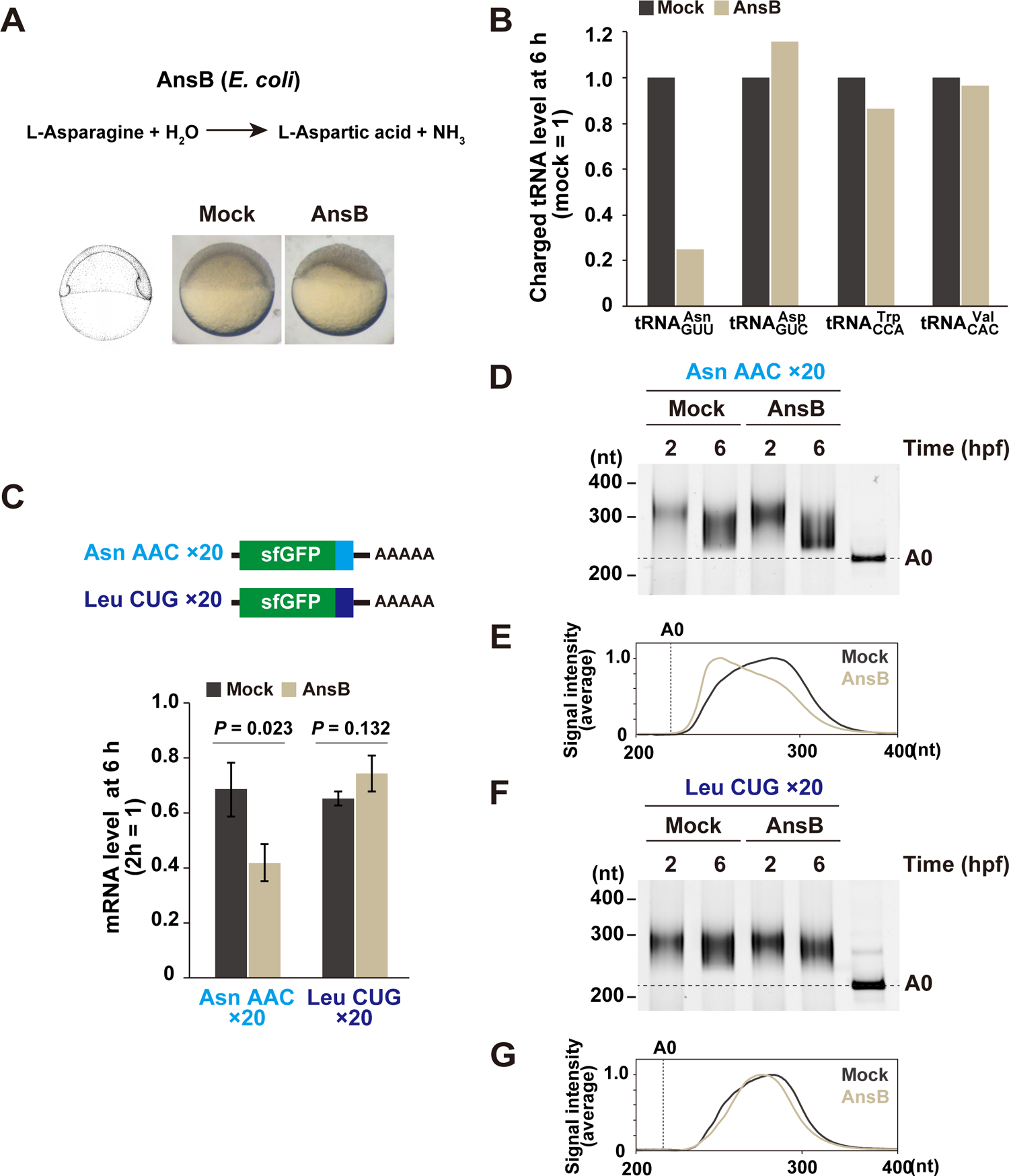
Modulation of charged tRNA amount in zebrafish embryos. (A) AnsB causes asparagine deprivation by converting asparagine to aspartic acid. Mock-injected and AnsB-overexpressing zebrafish embryos at 6 hpf are shown. (B) Charged tRNA levels in mock-injected and AnsB-overexpressing embryos at 6 hpf. Total and charged tRNAs were measured by qRT-PCR coupled to the OXOPAP assay. The relative charged tRNA level in mock-injected embryos was set to one. (C) qRT-PCR analysis of Asn AAC and Leu CUG codon tag reporter mRNAs at 6 hpf relative to 2 hpf in mock-injected and AnsB-overexpressing embryos. Experiments were repeated three times, and the average is shown. Error bars show SD. P-values were calculated using two-sided Student’s t-test. (D) Poly(A) tail analysis of Asn AAC codon tag reporter mRNA at 2 and 6 hpf. The developmental stages are shown above as hpf. The lane labeled A0 shows the 3’ UTR fragment without a poly(A) tail. (E)) Quantification of the PAT assay in (D). (F) Poly(A) tail analysis of Leu CUG codon tag reporter mRNA at 2 and 6 hpf. (G) Quantification of the PAT assay in (F).

### Codon-mediated decay occurs independent of Znf598

In theory, the slowed ribosome at the decoding step may drive other types of cotranslational mRNA decay such as NGD. Thus, we tested whether the codon effects observed in our PACE experiments represented codon-mediated decay or a mixed output with NGD.

As NGD has not been previously reported in zebrafish, we first had to investigate whether the ribosome stall induces NGD in zebrafish embryos. For this purpose, we focused on the ribosome stalling sequence from human cytomegalovirus (hCMV) uORF2, whose peptide arrests the ribosome on the stop codon together with eRF1 (Matheisl et al., 2015). Here, we hypothesized that hCMV uORF2 causes ribosome stalls to be strong enough to induce NGD in zebrafish embryos. When we inserted codon-optimized hCMV uORF2 at the 3’ end of the sfGFP ORF, protein output was significantly reduced in zebrafish embryos (Figure 5A and 5B). As reported, mutating the diproline motif at the end of uORF2, critical amino acids for ribosome stalling, partially restored the protein output (Matheisl et al., 2015). We then measured the stability of these reporter mRNAs and found that hCMV uORF2 reduced mRNA stability in a diproline motif-dependent manner (Figure 5C). Notably, the PAT assay detected no significant deadenylation of the reporter mRNAs (Figure 5D and E). These results suggested that hCMV uORF2 induced mRNA decay in a deadenylation-independent manner, distinct from codon-mediated decay.

**Figure 5.**
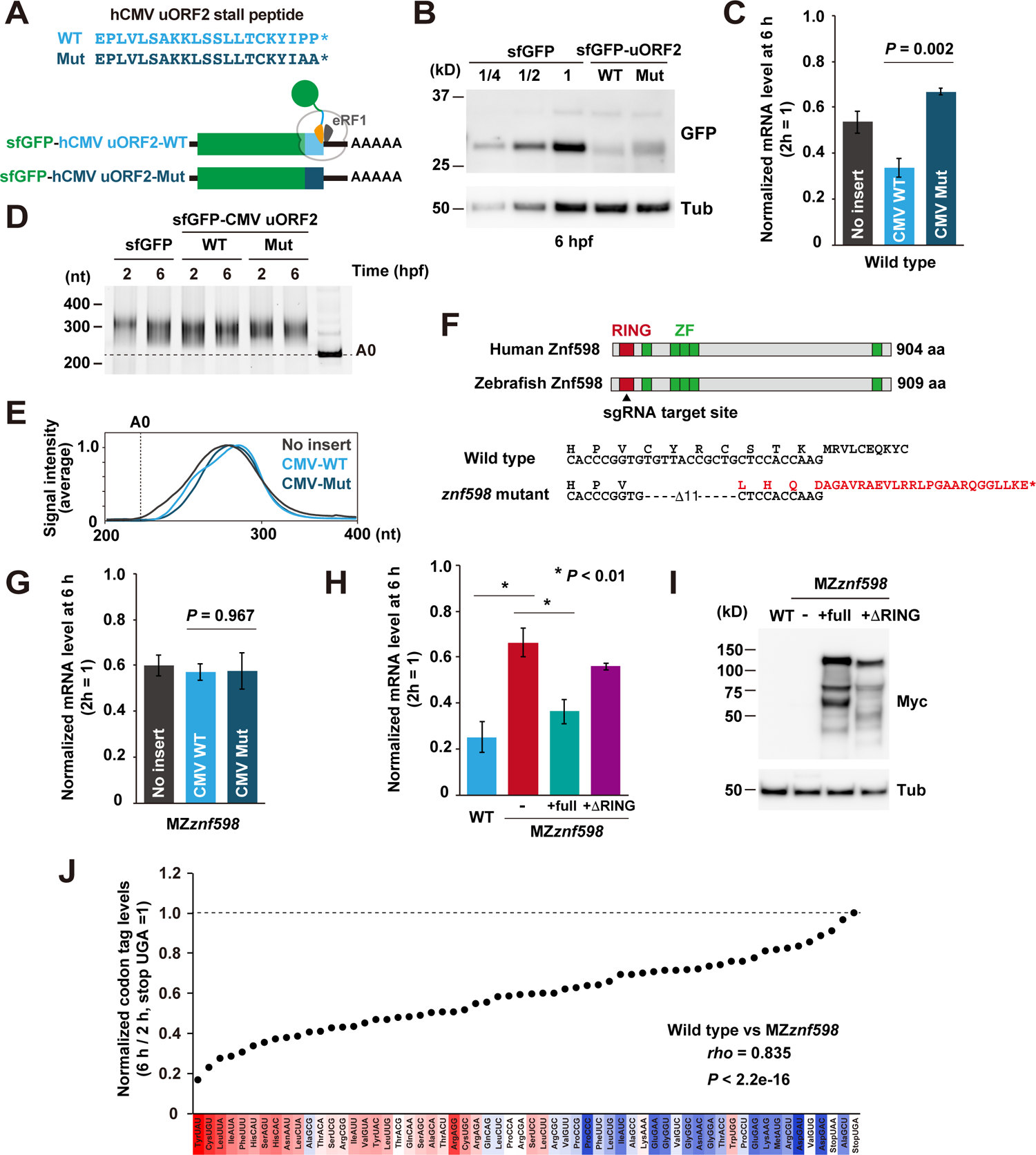
The effect of ribosome stall on mRNA stability. (A) A scheme of the hCMV uORF2 reporter mRNAs. Wild-type hCMV uORF2 (light blue) or its mutant (dark blue) was inserted at the end of the superfolder GFP (sfGFP) ORF. (B) Western blotting to detect the sfGFP protein levels at 6 hpf. Tubulin (Tub) was detected as a loading control. (C) qRT-PCR analysis of hCMV uORF2 reporter mRNA levels in wild-type embryos at 6 hpf relative to 2 hpf. The experiments were repeated three times. Error bars show SD. (D) Poly(A) tail analysis of hCMV uORF2 reporter mRNAs at 2 and 6 hpf. The developmental stages are shown above as hpf. The lane labeled A0 shows the 3’ UTR fragment without a poly(A) tail. (E)) Quantification of the PAT assay in (D). (F) Scheme of human and zebrafish Znf598 proteins. The sgRNA target site is indicated with an arrowhead. Genome sequences of wild-type and *znf598* mutant zebrafish are shown below. The predicted amino acid sequence of mutated Znf598 is shown in red. The asterisk indicates the premature stop codon. (G) qRT-PCR analysis of hCMV uORF2 reporter mRNA levels in MZ*znf598* embryos at 6 hpf relative to 2 hpf. The experiments were repeated three times. Error bars show SD. (H) qRT-PCR analysis of hCMV uORF2 reporter mRNA levels in wild-type embryos, MZ*znf598* embryos and MZ*znf598* embryos injected with an mRNA encoding Myc-tagged full-length (full) or RING domain deleted (*Δ*RING) Znf598. mRNA levels at 6 hpf relative to 2 hpf are shown. The experiments were repeated three times. Error bars show SD. (I) Western blotting to detect the Myc-tagged Znf598 proteins at 6 hpf. Tubulin was detected as a loading control. (J) Results of PACE in MZ*znf598* zebrafish embryos. Black circles show the relative stability of the reporter mRNAs. The stability of a codon-tag reporter with a UGA stop codon is set to one. Averages of two biological replicates are shown. The relative effect of each codon on mRNA stability in wild-type embryos measured in Figure 2B is shown as a color gradation with red (strong) to blue (weak). In C, G, and H, P-values were calculated using two-sided Student’s t-test.

RQC-coupled NGD in yeast requires the E3 ubiquitin ligase Hel2 and subsequent ubiquitination of ribosomal protein uS10 (Ikeuchi et al., 2019). To characterize mRNA decay induced by hCMV uORF2, we generated a zebrafish mutant for Znf598, a vertebrate homolog of Hel2, by using CRISPR-Cas9. We obtained a strain containing an 11 bp deletion in exon 1 of the *znf598* locus, which was predicted to induce a frameshift and produce a protein product truncated within the RING finger domain (Figure 5F). Embryos lacking both maternal and zygotic Znf598 (MZ*znf598*) were obtained by crossing homozygous *znf598* mutants. *znf598* mRNA was significantly reduced in MZ*znf598* (Figure S3A), likely reflecting the premature termination of translation followed by nonsense-mediated mRNA decay (Maquat, 2004). The detailed phenotype of our *znf598* mutant will be described elsewhere. Analysis of the reporter mRNA stability revealed that hCMV uORF2 did not induce mRNA decay in MZ*znf598* embryos (Figure 5G). Rescue experiments revealed that the mRNA decay activity of hCMV uORF2 was restored by full-length Znf598 but not by a deletion construct lacking the RING domain (Figure 5H and I). These results showed that specific ribosome stalls induced mRNA decay in a Znf598-dependent manner in zebrafish embryos.

This observation led us to test the impact of Znf598 on codon effects measured by PACE (Figure 5J). Overall, we found that most codon effects were maintained in MZ*znf598* compared to wild type, with a high correspondence (the correlation coefficients 0.826 [Pearson] and 0.835 [Spearman]) (Figure 5J and S3B). However, we noted that a few codons changed their effects in MZ*znf598*: Ala GCG turned into a destabilizing codon, whereas Lys AAA and Trp UGG codon tag reporter mRNAs were more stable in MZ*znf598* than wild type. Since ribosome stalls formed by consecutive AAA and UGG codons are subject to RQC in yeast and humans (Dimitrova et al., 2009; Juszkiewicz and Hegde, 2017; Mizuno et al., 2021; Sundaramoorthy et al., 2017), these codon tags might induce mRNA decay in part via the NGD-like pathway in zebrafish embryos. We draw two conclusions from these results: 1) Znf598-dependent NGD-like activity exists in zebrafish embryos and 2) most if not all of the codon effects measured by PACE are distinct from Znf598-dependent mRNA decay.

### Codon-mediated decay occurs during productive translation

To further characterize the nature of the ribosome slowdown that leads to codon-mediated decay, we performed a tandem ORF reporter assay that could quantitatively detect ribosome stalling in mammalian cells (Juszkiewicz and Hegde, 2017; Sundaramoorthy et al., 2017). Myc-tagged enhanced GFP (EGFP) and HA-tagged DsRed-Express (DsRedEx) were separated by two porcine teschovirus-1 2A (P2A) sequences that cause skipping of peptide bond formation (Kim et al., 2011), and a sequence to be tested for the efficiency of translation elongation was inserted between the two P2A sequences (Figure 6A). If the inserted sequence inhibited the elongation process of the ribosome, the DsRedEx/EGFP ratio would be reduced in this experimental setup. In vitro synthesized mRNA was injected into 1-cell stage zebrafish embryos, and the protein expression was analyzed at 3 hpf by western blotting to detect protein levels before the onset of codon-mediated decay.

**Figure 6.**
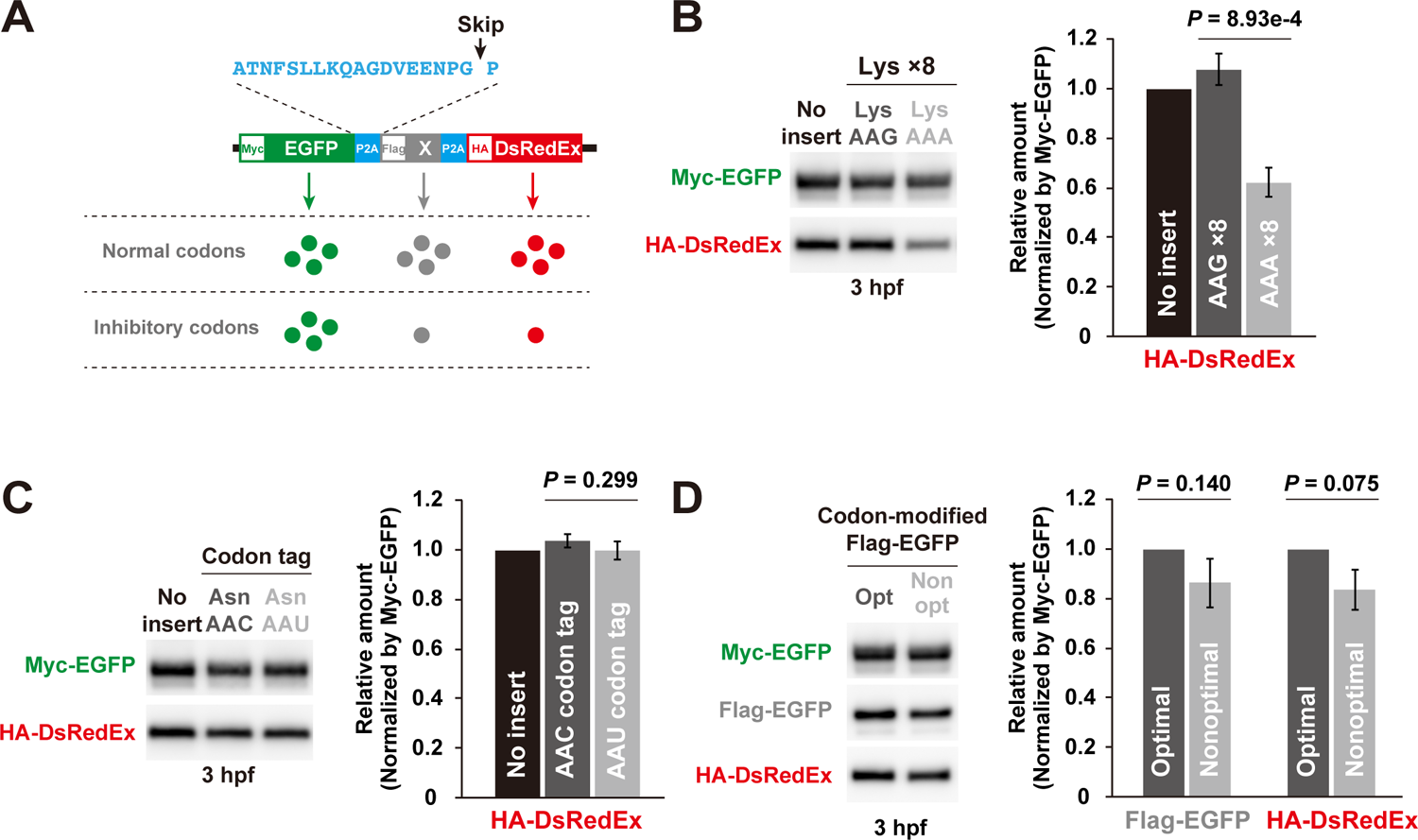
Analysis of the translation elongation rate in zebrafish embryos. (A) A schematic of the tandem ORF assay. Myc-tagged EGFP ORF (green) and HA-tagged DsRedEx ORF (red) are separated by two P2A translation skipping sequences (light blue). A sequence to be tested (gray) is inserted between the two P2A sequences. (B) Results of the tandem ORF assay with Lys AAG×8 and Lys AAA×8. Representative western blotting detecting Myc-tagged EGFP and HA-tagged DsRedEx at 3 hpf is shown on the left. Quantification of the HA-DsRedEx levels normalized by Myc-EGFP is shown on the right. The HA-DsRedEx level with no insert sequence is set to one. (C) Results of the tandem ORF assay with Asn AAC and Asn AAU codon tags. Representative western blotting detecting Myc-tagged EGFP and HA-tagged DsRedEx at 3 hpf is shown on the left. Quantification of the HA-DsRedEx levels normalized by Myc-EGFP is shown on the right. The HA-DsRedEx level with no insert sequence is set to one. (D) Results of the tandem ORF assay using EGFP ORFs with different codon optimality. Representative western blotting detecting Myc-tagged EGFP, Flag-tagged codon-modified EGFP and HA-tagged DsRedEx at 3 hpf is shown on the left. Quantification of Flag-EGFP and HA-DsRedEx levels normalized by Myc-EGFP is shown on the right. The Flag-EGFP or HA-DsRedEx level with no insert sequence was set to one. All experiments were repeated three times, and the average is shown. Error bars show SD. P-values were calculated using two-sided Student’s t-test.

To verify the sensitivity of the assay, we first analyzed the effect of the consecutive Lys AAA codons that are known to induce ribosome stall, collision, and RQC in other organisms (Ito-Harashima et al., 2007; Juszkiewicz and Hegde, 2017; Sundaramoorthy et al., 2017). Consistent with previous studies, ribosome stall at the eight consecutive AAA codons was detected as an approximately 40 % reduction in DsRedEx expression compared to the consecutive Lys AAG codons (Figure 6B). We then analyzed the effect of two synonymous AAC and AAU Asn codon tags, which showed opposite impacts on mRNA stability in PACE (Figure 2B and C). We found that Asn codon tag reporters showed almost identical DsRedEx/EGFP ratios (Figure 6C). This result suggested that ribosomes were not significantly retarded on the destabilizing codons. To confirm this idea with other codons, we compared two EGFP ORFs encoded by either optimal or nonoptimal codons throughout the ORF (Mishima and Tomari, 2016). The inserted nonoptimal EGFP ORF tended to reduce downstream DsRedEx expression, but the difference was quite subtle and statistically insignificant (Figure 6D). Hence, the translation of >200 nonoptimal codons was still more efficient than that of a short, stall-prone polyA stretch. These results showed that the ribosome slowdown required for inducing codon-mediated decay is far more transient than the ribosome stall required for inducing ribosome collision and RQC.

## Discussion

In this study, we developed a reporter-based assay named PACE to analyze codon-mediated decay in zebrafish embryos. PACE was sensitive enough to detect translation-dependent, codon-specific effects on mRNA stability in zebrafish embryos (Figure 2). PACE relies on relatively simple bioinformatics analysis and is potentially applicable to other organisms capable of undergoing mRNA injection or transfection. Optimization of the codon tag sequence and the inclusion of the four codon tags missed during its construction or RNA-seq will broaden the utility of PACE.

Previous studies estimated codon effects by calculating the correlation between the codon frequency and mRNA stability for thousands of endogenous mRNAs (Bazzini et al., 2016; Forrest et al., 2020; Harigaya and Parker, 2016; Mishima and Tomari, 2016; Presnyak et al., 2015). These analyses took advantage of averaging codon effects in multiple different positions on the transcriptome. However, the correlation tended to be modest in part due to the presence of factors that affect mRNA stability independent of codons (e.g., RNA-binding proteins and microRNAs) (Medina-Muñoz et al., 2021). The ORFome approach minimizes this disadvantage by fixing the 5’ and 3’ UTR sequences, yet it also depends on a correlation study, and its target is limited to endogenous mRNA sequences that are under evolutionary constraints (Bazzini et al., 2016; Narula et al., 2019; Wu et al., 2019). In contrast, PACE places each codon in a single artificial context to directly evaluate the codon effects on mRNA stability. As the findings of PACE in this study were largely consistent with those of CSC, our study provides additional, experimental validation of previously reported codon effects on mRNA stability in zebrafish (Figure 2B-E and S1D). Furthermore, we showed several lines of evidence indicating that codon effects stem from the decoding process in zebrafish. First, codons with destabilizing effects had fewer corresponding tRNA amounts than those with stabilizing effects (Figure 2D). Second, the occupancy of the ribosome with an A-site codon for a polar amino acid was negatively correlated with the codon’s effect on mRNA stability (Figure 3C) and the corresponding tRNA amount (Figure 3G). Third, experimental reduction of Asn tRNA availability conferred a destabilizing effect to the Asn AAC codon (Figure 4). Together, our study supports the view wherein a slow ribosome waiting for decoding by tRNAs triggers mRNA degradation in eukaryotes.

One limitation of our analysis was that a correlation between the codon effect and the decoding process was not observed with codons for nonpolar and charged codons (Figure 3D and 3E). Although the exact reason is unknown, we noted that the ribosome occupancy and tRNA amounts were poorly correlated with these codons (Figure 3H and I). As either local or net hydrophobicity and the charge of the nascent peptides affects ribosome movement and functionality (Chadani et al., 2017, 2016; Charneski and Hurst, 2013; Dao Duc and Song, 2018; Lu and Deutsch, 2008; Weinberg et al., 2016), one possibility is that ribosome occupancy on these codon tags predominantly reflects the effect of nascent peptides; therefore, a precise measurement of the ribosome waiting for decoding is hampered. Future studies are necessary to confirm whether the decoding process of codons for nonpolar and charged amino acids also serves as a determinant of mRNA stability in zebrafish.

Recent studies in yeast have provided mechanistic insights into codon-mediated decay and NGD, establishing them as distinct cotranslational mRNA decay pathways (Buschauer et al., 2020; D’Orazio et al., 2019; Ikeuchi et al., 2019; Webster et al., 2018). However, their definition is less clear in other organisms due to the lack of a connection between cis-elements and trans-acting factors. We showed that ribosome stall formed by hCMV uORF2 induced mRNA decay in a deadenylation-independent and Znf598-dependent manner, thus providing the first example of NGD in zebrafish embryos (Figure 5A-I). Irrespective of that, we found that the codon effects were largely maintained even in the absence of Znf598 (Figure 5J). Hence, cellular mechanisms clearly distinguish ribosome slowdown during the decoding process from ribosome stall by a problematic RNA or peptide and promote specific downstream mRNA decay pathways in zebrafish.

How does the cell distinguish ribosomes for codon-mediated decay and those for NGD? Although earlier yeast studies provided the insights, it is unclear whether the underlying molecular mechanisms for recognizing slow ribosomes and stalled ribosomes are evolutionally conserved in higher eukaryotes. Whereas Not5 is not conserved in vertebrates, Not3 shares similarity with Not5 in the N-terminal domain required for E-site binding. It is therefore possible that the yeast Not3 ortholog CNOT3 anchors the CCR4-NOT complex to the slow ribosome via the E site in vertebrates (Buschauer et al., 2020; Collart and Panasenko, 2017). The ubiquitination site(s) on the ribosome and downstream NGD effectors are also conserved in vertebrates (D’Orazio et al., 2019; Garzia et al., 2017; Ikeuchi et al., 2019; Juszkiewicz et al., 2020, 2018; Juszkiewicz and Hegde, 2017; Matsuo et al., 2017; Sundaramoorthy et al., 2017; Weber et al., 2020). Future studies will reveal the conservation and biological function of codon-mediated decay and NGD. As exemplified by our analysis with the *znf598* mutant, PACE will be a valuable tool to compare codon effects in multiple mutant backgrounds in such studies.

Although the ribosome slows down locally on a destabilizing codon, our tandem ORF assay showed that such slowdown had a limited impact on the overall translation elongation rate (Figure 6). This assay was sensitive enough to detect the inhibitory effect of RQC-prone consecutive AAA codons. Therefore, we propose that ribosome slowdown during codon-mediated decay does not reach a nonproductive stall and maintains a productive translation cycle. This model is reasonable given that codon-mediated decay regulates the stability of thousands of endogenous protein-coding mRNAs for essential metabolic genes and maternal genes (Bazzini et al., 2016; Forrest et al., 2020; Mishima and Tomari, 2016; Presnyak et al., 2015). One remaining question in codon-mediated decay is how such transient slowness of the ribosome leads to robust deadenylation and mRNA degradation. In the case of RQC/NGD, a prolonged and robust ribosome stall causes a collision with the following ribosome, which becomes a substrate for Hel2/Znf598 (Ikeuchi et al., 2019; Juszkiewicz et al., 2018; Matsuo et al., 2020). While the collision itself is estimated to occur when the elongation rate decreases several-fold (∼6 codons/second to ∼1 codon/second) (Juszkiewicz et al., 2018), live imaging of RQC revealed that a much longer stall is required for RQC to occur (∼0.12 codons/second) (Goldman et al., 2021). Our study indicated that ribosome slowdown required for triggering codon-mediated decay did not reach such a prolonged ribosome stall (Figure 6). Among the two deadenylases in the Ccr4-Not complex, Caf1 is involved in deadenylation related to codon optimality in yeast (Webster et al., 2018). As Caf1 is a processive deadenylase, only a transient interaction between the Ccr4-Not complex and a slow ribosome, possibly mediated by E-site recognition, may be sufficient to initiate continuous deadenylation (Viswanathan et al., 2004). Alternatively, ubiquitination of eS7 by Not4 acts as an additional scaffold on the ribosome to stably recruit the Ccr4-Not complex after transient slowdown (Buschauer et al., 2020). Kinetical analysis of codon-mediated decay with a defined reporter mRNA is required to understand its detailed mechanism of action.

## Materials and Methods

### Zebrafish

The zebrafish AB strain was raised and maintained at 28.5°C under standard laboratory conditions according to the Animal Experiment Protocol (2018-46) at Kyoto Sangyo University. Fertilized eggs were obtained by natural breeding. Embryos were developed in system water at 28.5°C. Bright field images were acquired with a SteREO Lumar V12 microscope and an AxioCam MRc camera with ZEN software (Zeiss, Jena, Germany) using spline mode.

### Microinjection

To synthesize mRNA for microinjection, RNA was transcribed from a linearized plasmid DNA template using the SP6-Scribe Standard RNA IVT Kit (CELLSCRIPT) and purified with the RNeasy Mini Kit (QIAGEN). The m^7^G cap structure was added to the purified RNA using the ScriptCap m^7^G Capping System (CELLSCRIPT). Capped mRNA was purified with the RNeasy Mini Kit. For sfGFP reporter mRNAs, the poly(A) tail was added using the A-Plus Poly(A) Polymerase Tailing Kit (CELLSCRIPT), and the mRNA was then purified with the RNeasy Mini Kit. Purified mRNAs were diluted with water to the following concentrations: sfGFP reporter mRNAs and PACE reporter library 50 ng/μl; MT-Znf598 mRNAs 100 ng/μl; AnsB 50 ng/μl; tandem ORF reporter mRNAs 100 ng/μl. GFP MO was injected as described previously (Mishima and Tomari, 2016). Microinjection was performed using an IM300 Microinjector (NARISHIGE). Approximately 1,000 pl of solution was injected per embryo within 15 min after fertilization.

### Generation of a *znf598* mutant strain by CRISPR-Cas9

sgRNA targeting exon 1 of zebrafish *znf598* (ENSDARG00000014945) was designed with CRISPRscan (Moreno-Mateos et al., 2015). The DNA template for sgRNA synthesis was prepared by PCR using gene-specific primer and sgRNA tail primer. sgRNAs were transcribed in vitro using T7 RNA Polymerase (TaKaRa) and purified with ProbeQuant G-50 microcolumns (Cytiva). m^7^G-capped nCas9 mRNA was prepared with the SP6-Scribe Standard RNA IVT Kit and ScriptCap m^7^G Capping System (CELLSCRIPT) using pCS2+nCas9n as a template (Jao et al., 2013). sgRNA and nCas9 mRNA were mixed at 50 ng/μl each and injected into fertilized eggs. Embryos were raised to adulthood, and the fishes were crossed with AB fish. F1 embryos were screened by PCR analysis using Multina (SHIMADZU). DNA sequences were determined by cloning PCR fragments and Sanger sequencing. We identified a strain with an 11 bp deletion in exon 1, as shown in Figure 5F. Siblings of F1 fishes were raised to adulthood, and the fish harboring a heterozygous mutation were identified by fin-clipping and genotyping PCR. The F1 fish were crossed with AB to dilute any nonspecific mutations, and the F2 fish were used to obtain homozygous mutants. Primers are listed in Table S1

### Plasmid construction

To construct reporter plasmids, pCS2+ (Rupp et al., 1994) was modified by PCR to generate pCS2+neo using primers y683fw, y683rv, y684fw and y684rv. The *suv39h1a* 3’ UTR was excised from the previously described EGFP reporter plasmid (Mishima and Tomari, 2016) and cloned into pCS2+neo by XhoI and XbaI. The amino acid sequence of the sfGFP ORF was converted to nucleotide sequences comprising the most frequent synonymous codons in *E. coli* using EMBOSS Backtranseq (http://www.ebi.ac.uk/Tools/st/emboss_backtranseq/). The first 25 nucleotides of the sfGFP ORF were identical to the EGFP ORF used in a previous study to maintain the target site for GFP Morpholino oligo (Mishima and Tomari, 2016). EcoRI and EcoRV sites were inserted before the stop codon. The sfGFP DNA fragment was synthesized by GeneArt Strings DNA Fragments service (Life Technologies) and cloned into the pCS2+neo-*suv39h1a* 3’ UTR by NcoI and XhoI. DNA oligonucleotides encoding hCMV uORF2 were annealed and cloned into the pCS2+neo-sfGFP-*suv39h1a* 3’ UTR by EcoRI and EcoRV.

To construct the tandem ORF reporter plasmid, the EGFP ORF was amplified by PCR using primers y670 and y671 and then amplified by y671 and y674. DsRedEx was amplified by PCR using y672 and y673 and then amplified by y675 and y673. The resulting EGFP fragment contained EGFP-P2A-Flag and the DsRedEx fragment contained P2A-HA-DsRedEx. Finally, the two fragments were joined by PCR using y670 and y673, generating EGFP-P2A-Flag-P2A-HA-DsRedEx. The final product was cloned into pCS2+MT containing the *suv39 h1a* 3’UTR using NcoI and XhoI sites. To insert the Lys AAG ×8 or AAA ×8 sequence, primers y713fw and y713rv or y714fw and y714rv were annealed and cloned downstream of Flag by EcoRV and EcoRI. Asn codontags were also cloned downstream of Flag by EcoRV and EcoRI. Optimal and nonoptimal EGFP described previously (Mishima and Tomari, 2016) were amplified by PCR using y707 and y708 and cloned downstream of Flag by EcoRV and EcoRI.

Full-length and *Δ*RING Znf598 ORF fragments were amplified by RT-PCR using primers y624 and y775 and cloned into pCS2+MT by XhoI and XbaI. AnsB was amplified by PCR from the genomic DNA of *E.coli* DH5*α* using primers y603 and cloned into pCS2+SBP-HA (Mishima and Tomari, 2016) by EcoRI and XhoI. Primers are listed in Table S1.

### PACE library

Oligonucleotides encoding codon tag sequences for all codons were synthesized as 64 pairs of single-strand DNA fragments. Complementary oligonucleotides were annealed and cloned into the pCS2+neo-sfGFP-*suv39h1a* 3’ UTR by EcoRI and EcoRV. The sequence of the cloned plasmids was confirmed by Sanger sequencing. Three codon tags, for UCU, UCA, and UAG, could not be cloned for unknown reasons. The cloned plasmids were mixed at an equimolar concentration and linearized by NotI. Capped and polyadenylated mRNAs were prepared using the mixed templates and purified as described above. The homogenous length of the PACE library mRNAs was confirmed by electrophoresis. The codon tag sequences are listed in Table S2.

### RNA sequencing

Total RNA was extracted from 40–50 embryos injected with the PACE library mRNAs using TRI Reagent (Molecular Research Center) and purified using the RNeasy Mini Kit (QIAGEN). After RNA integrity was confirmed using an Agilent RNA 6000 Nano Chip (Agilent Technologies, USA), total RNA samples were treated with Ribo-Zero (Illumina) and used for RNA-seq library preparation using an Illumina TruSeq Stranded Total RNA Library Prep Kit (Illumina, USA) according to the manufacturer’s instructions with multiplexing. The pooled libraries were sequenced on HiSeq4000 or NextSeq500 (Illumina) sequencing platforms with single-end sequencing of 65 (HiSeq4000) or 76 bp (NextSeq500) lengths.

### Ribosome footprint profiling

Ribosome footprint profiling using zebrafish embryos was performed as described previously (Han et al., 2020; Mito et al., 2020). Briefly, 50–60 zebrafish embryos injected with PACE reporter mRNAs were snap-frozen at the sphere stage and lysed with lysis buffer containing cycloheximide at 100 μg/ml (Sigma-Aldrich). An S-400 HR gel filtration spin column (GE Healthcare) was used to isolate the ribosomes. RNA fragments ranging from 26-34 nt were selected for monosome profiling. PCR-amplified DNA libraries were gel-excised and sequenced with the HiSeq4000 platform (Illumina).

### Data analysis

RNA-seq reads for PACE analysis were analyzed using Galaxy on public servers (Goecks et al., 2010). Single-ended reads matched to the codon tag sequences were counted by Salmon quant (Patro et al., 2017) with default parameters using the sequences listed in Table S2 as references. The number of reads on each codon tag was normalized to the total reads mapped to the zebrafish genome (GRCz10/danRer10) by TopHat v2.1.1. To calculate the stability of the codon tag reporter mRNAs, the normalized read numbers at 6 hpf were divided by the normalized read numbers at 2 hpf. The stability of each codon tag reporter mRNA was further normalized by the value of a UGA codon tag reporter mRNA. The results from two biological replicates were averaged and used for the analysis.

Sequences from the ribosome profiling data were aligned against ncRNA sequences (GRCz11/danRer11.noncoding RNA genes data set) and mitochondrial gene sequences using Bowtie2; sequences that aligned were removed. The remaining sequences were aligned to the codon tag reporter sequences or the zebrafish genome (GRCz11/danRer11) using TopHat. Specific A-site assignment was determined by sequences aligned to the zebrafish genome, calibrated from footprints mapped to the beginning of the CDSes. The raw footprint read count at the test codon or spacer codon (as described in Figure 3A) was counted for further calculations.

### qRT-PCR

qRT-PCR analysis of mRNAs was performed as described previously (Mishima and Tomari, 2016). Briefly, total RNA was prepared using TRI Reagent (Molecular Research Center) and cDNA was synthesized using the PrimeScript RT reagent kit with gDNA eraser (TaKaRa). A random hexamer was used for cDNA synthesis. qRT-PCR was performed using SYBR Premix Ex TaqII (Tli RNaseH Plus) and the Thermal Cycler Dice Real Time System (TaKaRa). The results were analyzed based on standard curves from a serial dilution of cDNA. The data were normalized using 18S rRNA as a reference, and the mRNA level at 6 hpf was normalized to the data at 2 hpf. The primer sequences are listed in Table S1.

### Measurement of the charged tRNA level

To measure total tRNA and the charged tRNAs, we modified the OXOPAP assay (Gaston et al., 2008) and combined it with qRT-PCR analysis. Total RNA was purified with TRI reagent and dissolved in storage buffer (50 mM NaOAc pH 5.2 and 1 mM EDTA). For charged tRNA, 1 μg of the total RNA was incubated in oxidization buffer (50 mM NaIO_4_ and 100 mM NaOAc pH 5.2) supplemented with RNasin (Promega) for 30 min at 25°C to oxidize the nonacylated tRNA 3’ end. For the total tRNA, 1 μg of the total RNA was incubated in control buffer (50 mM NaCl and 100 mM NaOAc pH 5.2) supplemented with RNasin for 30 min at 25°C. Reactions were quenched with 100 mM glucose, incubated for 5 min at 25°C, and purified with G-25 columns (Cytiva). RNA samples were deacylated by adding 3.4 μl of 1 M Tris-HCl pH 9.0 and incubated for 30 min at 37°C. RNA samples were purified by ethanol precipitation and dissolved in 6.5 μl of RNase-free water. Four microliters of the RNA sample was denatured at 65°C for 5 min, and polyadenylated by the Poly(A) Polymerase Tailing Kit (CELLSCRIPT) in a 10 μl reaction in the presence of 2 U ePAP at 37°C for 60 min. Two-point five microliters of the polyadenylated sample was used for cDNA synthesis by the PrimeScript RT reagent kit with gDNA eraser (TaKaRa), with the y625 primer in Table S1. Reverse transcription was performed at 50°C for 30 min.

qRT-PCR was performed as described above using the tRNA-specific forward primers shown in Table S1. Specific amplification was confirmed by dissociation curve analysis followed by gel electrophoresis and sequencing. The charged tRNA level relative to the total tRNA level was calculated by the ΔΔCt method (ΔCt of total tRNA sample – ΔCt of charged tRNA sample). The charged tRNA levels of the AnsB-overexpression experiment were normalized to those of the mock injection experiment.

### PAT assay

The PAT assay was performed as described previously (Mishima and Tomari, 2016). Briefly, 150 ng of total RNA was incubated with 75 U of yeast poly(A) polymerase (PAP) (Affymetrix) in the presence of GTP/ITP mix at 37°C for 60 min. cDNA was synthesized at 44°C for 15 min using the PrimeScript RT reagent kit with gDNA eraser (TaKaRa) and a y300 PAT universal C10 primer. PAT-PCR was performed using a 3’ UTR-specific forward primer and a y300 PAT universal C10 primer with GoTaq Green Master Mix (Promega). The primer sequences are provided in Table S1. The PCR products were separated by 6% PAGE in 0.5×TBE. Gels were stained with GelRed (Biotium) and the signals were detected using LAS4000 (GE Healthcare) or Amersham Imager 680 (GE Healthcare). Signals were quantified using the “Plot profiles” function of the ImageJ software (http://imagej.nih.gov/ij/).

### Western blotting

Proteins were detected using anti-Myc (MBL My3 mouse monoclonal, 1:4,000), anti-HA (MBL TANA2 mouse monoclonal 1:4,000), anti-FLAG (MBL FLA-1 mouse monoclonal 1:4,000), anti-GFP (MBL No. 598 rabbit polyclonal, 1:3000), and anti-*α*-tubulin-pAb HRP-DirecT (MBL PM054-7 rabbit polyclonal 1: 10,000) antibodies. The signals were developed using Luminata Forte (Millipore) and detected using LAS4000 (GE Healthcare) or Amersham Imager 680 (GE Healthcare). Signals were quantified using the “Measure” function of the ImageJ software (http://imagej.nih.gov/ij/).

### Sequence data

The accession number for the sequence data reported in this paper is GEO: GSE173179 and GSE173604.

## Acknowledgments

This work was supported by the Japan Society for the Promotion of Science (JSPS) (JP18H02370), the Ministry of Education, Culture, Sports, Science and Technology (MEXT) (JP17H05593, JP17H05662) the Japan Agency for Medical Research and Development (AMED) (AMED PRIME, JP21gm6310017), and the Inamori Foundation to YM. SI was supported by JSPS (JP17H04998 and JP19K22406), MEXT (JP20H05784), AMED (AMED-CREST, JP21gm1410001), RIKEN (Pioneering Project “Biology of Intracellular Environments” and Aging Project), and the Takeda Science Foundation. SK was supported by JSPS (JP21H02513) and MEXT-Supported Program for the Strategic Research Foundation at Private Universities (S1511023). PH was an International Program Associate of RIKEN. We thank Yukihide Tomari and Kaori Kiyokawa for their support during the initial phase of this project and Ariel Bazzini for sharing zebrafish CSC and tRNA sequencing data and engaging in discussions. We thank our laboratory members for discussions and their critical comments on the project, Kimi Wakabayashi for fish maintenance, Kaori Kaminoyama and Tomoaki Sakamoto for RNA-Seq, and Mari Mito for technical assistance. We thank Shirahide laboratory at the Institute for Quantitative Biology for RNA-Seq. DNA libraries for ribosome profiling were sequenced by the Vincent J. Coates Genomics Sequencing Laboratory at UC Berkeley, which is supported by the National Institutes for Health (NIH) Instrumentation Grant (S10 OD018174). Computational analysis was supported by the supercomputer HOKUSAI Sailing Ship in RIKEN.

## Author contributions

Y.M. conceived and designed the experiments. P.H. and S.I. analyzed the ribosome profiling data. S.K. provided RNA-Seq data. Y.M. and S.I. wrote the manuscript. All the authors discussed the results and approved the manuscript.

---

**Supplemental figure 1.**
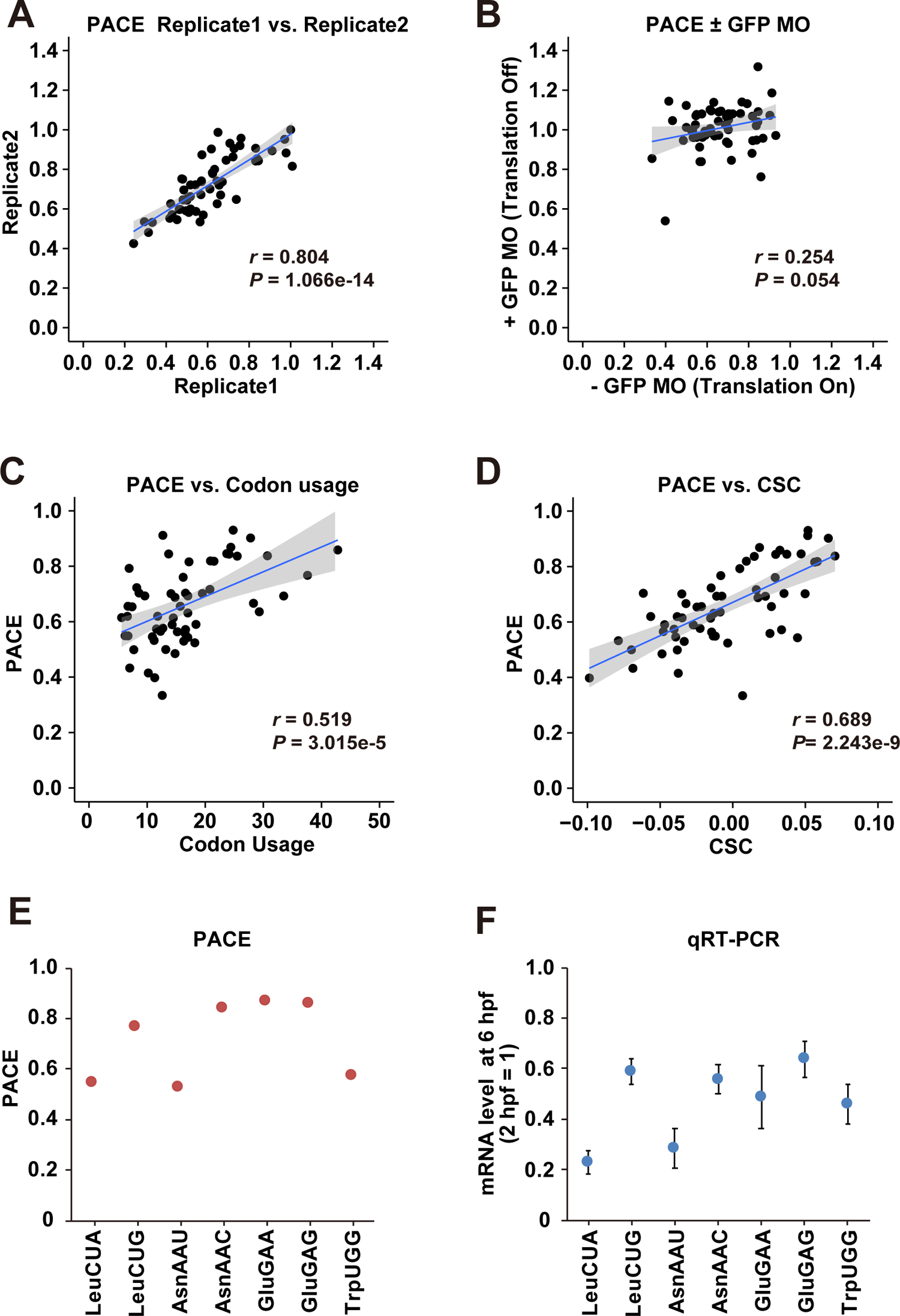
Validation of PACE, related to Figure 2. (A) A scatter plot showing the reproducibility of the codon effects measured by PACE in two replicates. (B) A scatter plot comparing the codon effects measured by PACE in the absence (x-axis) and presence of GFP MO (y-axis). (C) A scatter plot comparing codon usage in the zebrafish genome (per thousand codons, x-axis) and the codon effects measured by PACE (y-axis). (D) A scatter plot comparing CSCs (Bazzini et al., 2016, x-axis) and the codon effects measured by PACE (y-axis). (E and F) Comparison of the codon effects measured by PACE (E) with the results of qRT-PCR (F). mRNA levels at 6 hpf relative to 2 hpf are shown. The experiments were repeated three times. Error bars show SD. In A-D, the regression line is shown in blue and the 95% confidence interval is shown in gray. *r* represents Pearson’s correlation. The significance was calculated by Student’s t-test.

**Supplemental figure 2.**
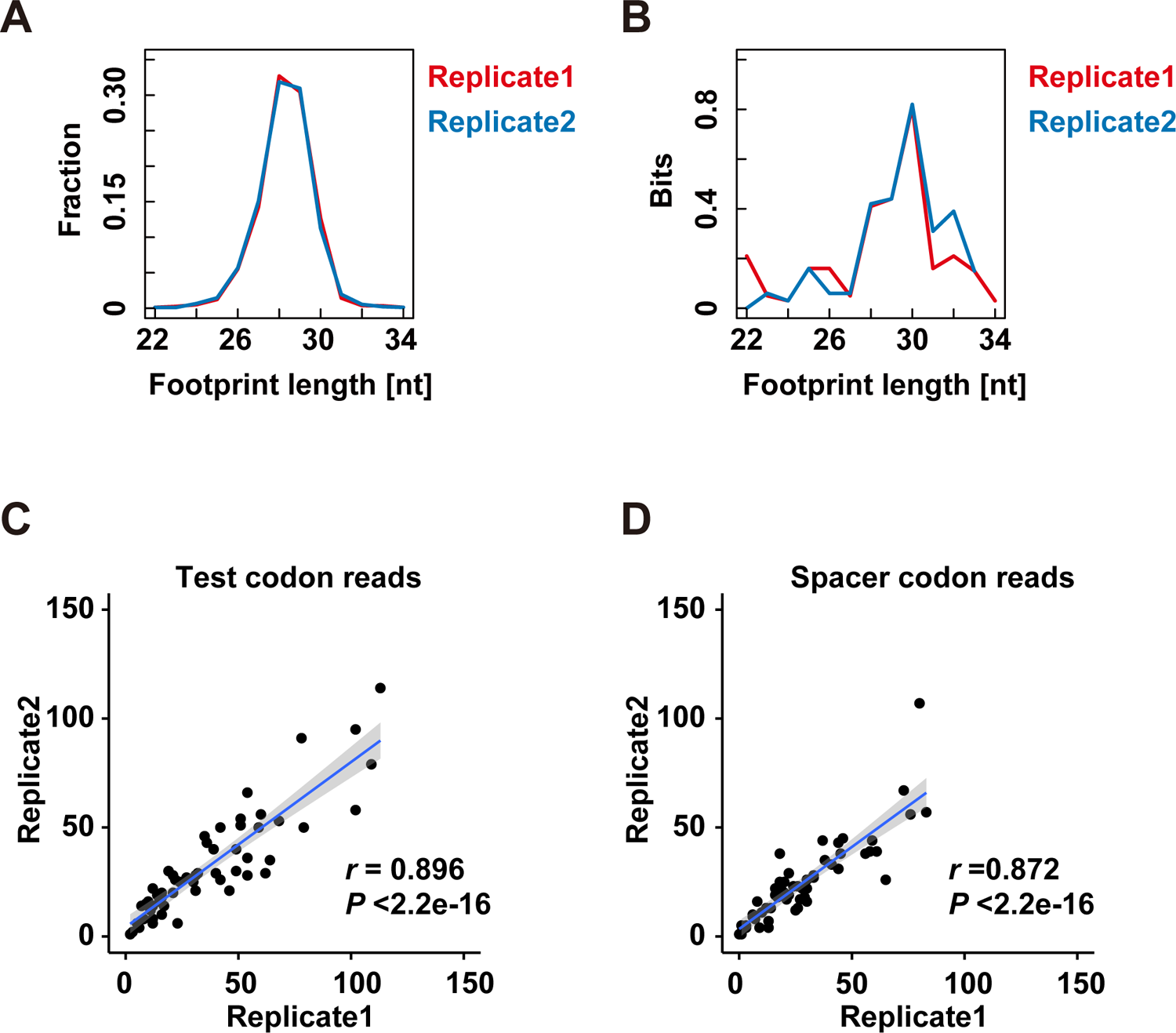
Validation of ribosome footprint analysis, related to Figure 3. (A) Distribution of the ribosome footprint lengths derived from codon-tag sequences. The results of the two replicates are plotted separately in red and blue. (B) Information content of each reading frame using ribosome footprints aligned to the codon tag reporter sequences. The results of the two replicates are plotted separately in red and blue. (C and D) Scatter plots showing the reproducibility of ribosome footprint analysis with PACE reporters. Two replicates for test codon reads (C) and spacer codon reads (D) are shown. The regression line is shown in blue and the 95% confidence interval is shown in gray. *r*, Pearson’s correlation. Significance was calculated by Student’s t-test.

**Supplemental figure 3.**
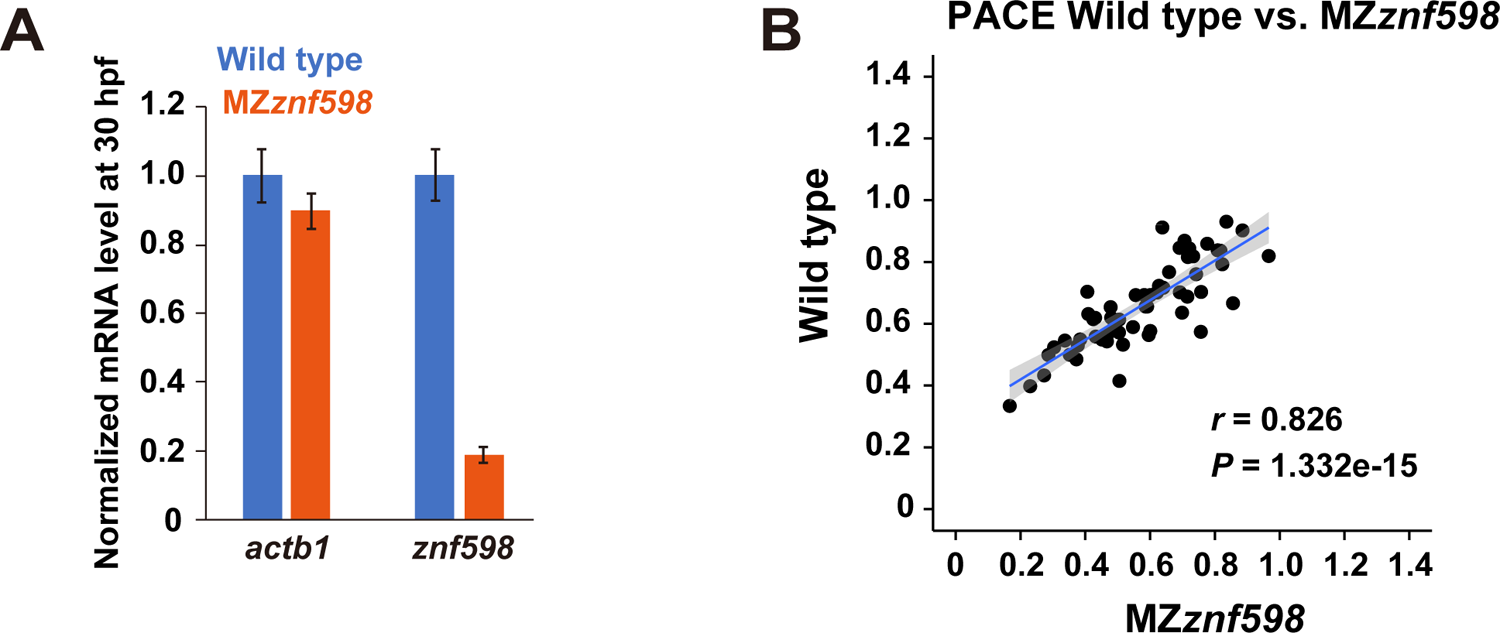
Validation of MZ*znf598* mutant, related to Figure 5. (A) qRT-PCR analysis of *actb1* and *znf598* mRNA levels in wild-type and MZ*znf598* embryos at 24 hpf. The results are normalized with the 18S rRNA level. The mRNA level in wild type is set to one. The experiments were repeated three times. Error bars show SD. (B) A scatter plot comparing the codon effects measured by PACE in MZ*znf598* (x-axis) and wild-type (y-axis) embryos. The regression line is shown in blue and the 95% confidence interval is shown in gray. *r*, Pearson’s correlation. Significance was calculated by Student’s t-test.

**Table S1.**
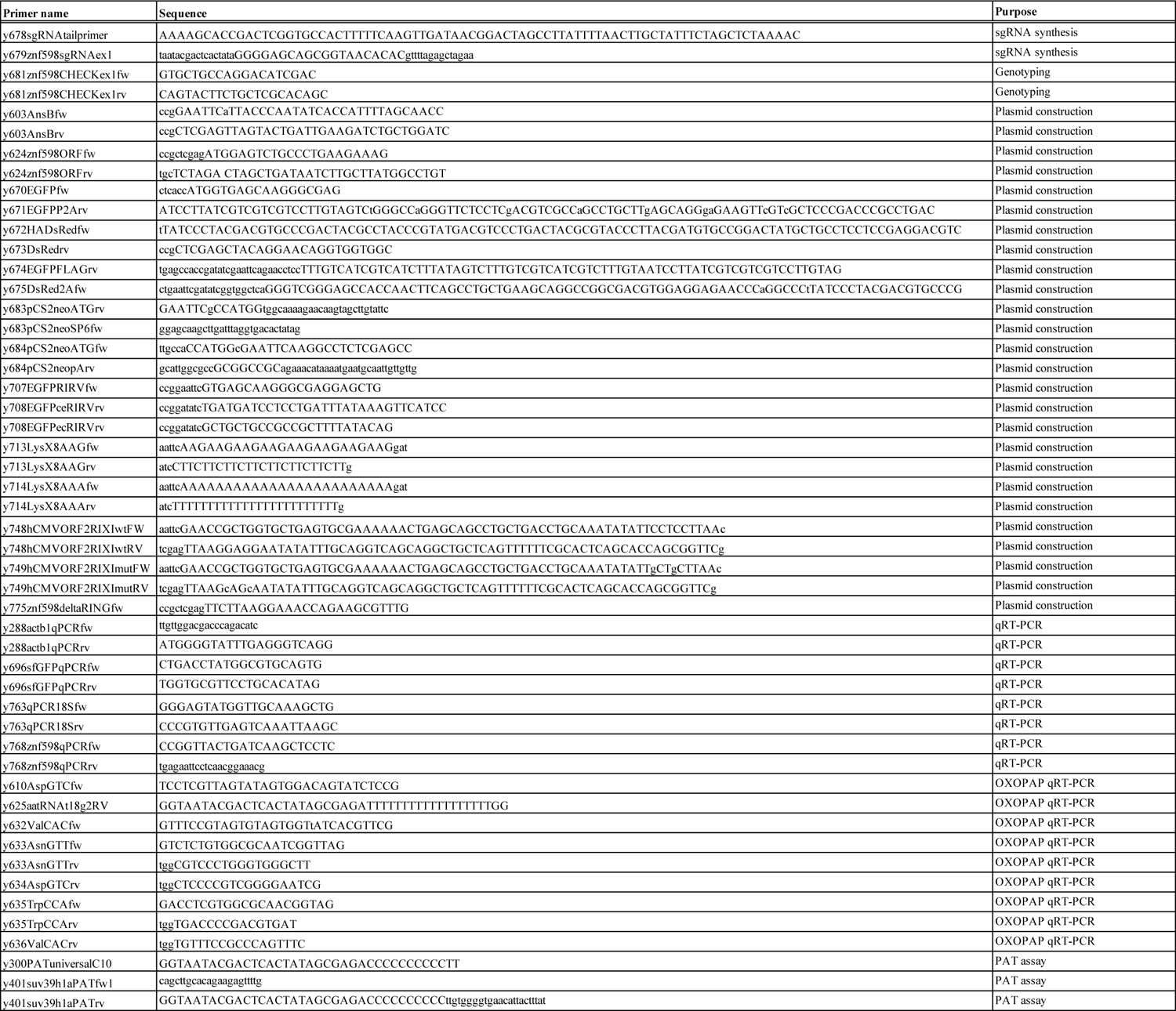
List of primers used in this study

**Table S2.**
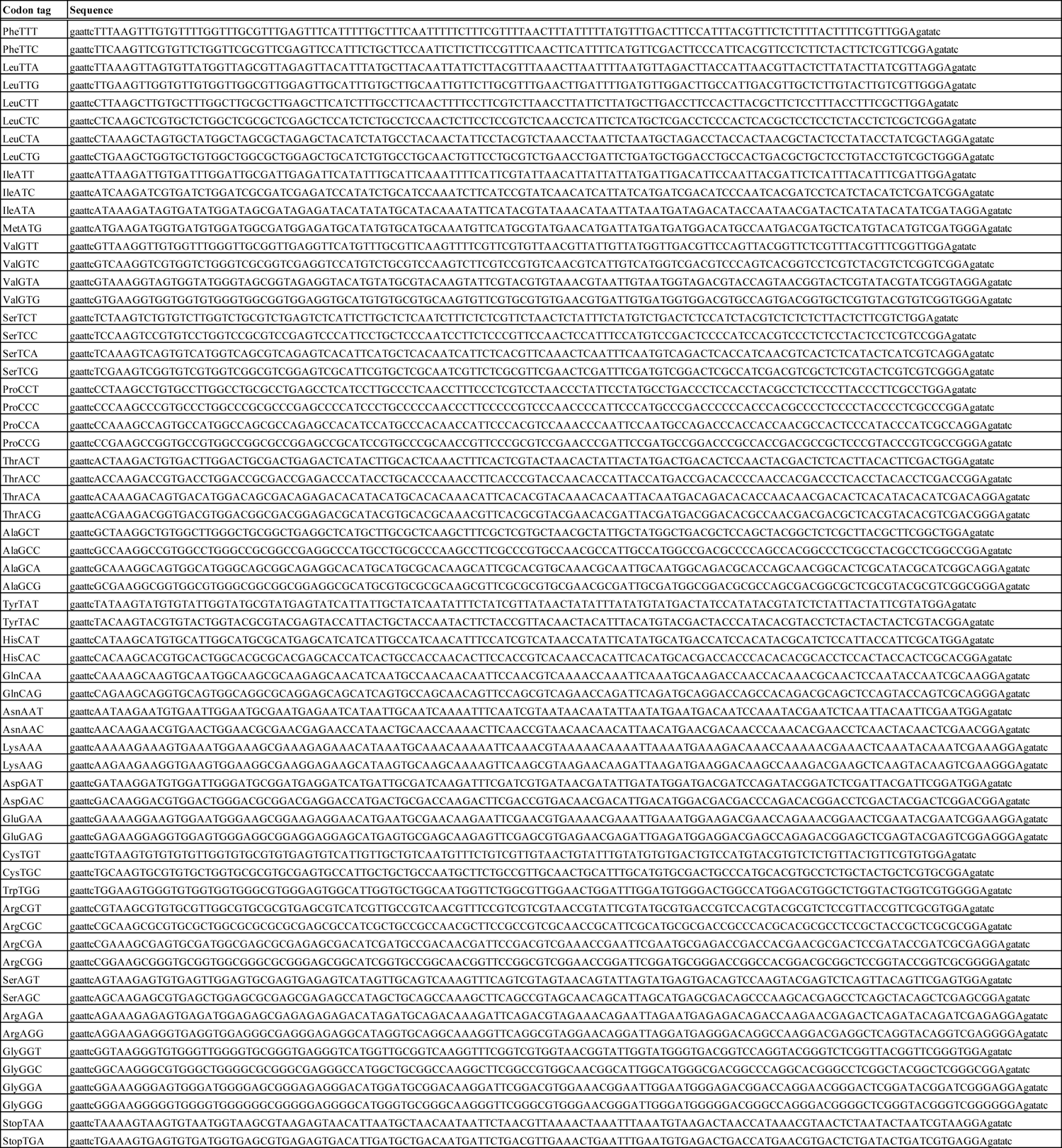
**List of codon tag sequences used in this study**

